# Low Repeatability of Aversive Learning in Zebrafish (*Danio rerio*)

**DOI:** 10.1101/2020.11.16.385930

**Authors:** Dominic Mason, Susanne Zajitschek, Hamza Anwer, Rose E O’Dea, Daniel Hesselson, Shinichi Nakagawa

## Abstract

Aversive learning – avoiding certain situations based on negative experiences – can profoundly increase fitness in animal species. The extent to which this cognitive mechanism could evolve depends upon individual differences in aversive learning being stable through time, and heritable across generations, yet no published study has quantified the stability of individual differences in aversive learning using the repeatability statistic, *R* (also known as the intra-class correlation). We assessed the repeatability of aversive learning by conditioning approximately 100 zebrafish *(Danio rerio)* to avoid a colour cue associated with a mild electric shock. Across eight different colour conditions zebrafish did not show consistent individual differences in aversive learning (*R* = 0.04). Within conditions, when zebrafish were twice conditioned to the same colour, blue conditioning was more repeatable than green conditioning (*R* = 0.15 and *R* = 0.02). In contrast to the low repeatability estimates for aversive learning, zebrafish showed moderately consistent individual differences in colour preference during the baseline period (i.e. prior to aversive conditioning; *R* ~ 0.45). Overall, aversive learning responses of zebrafish were weak and variable (difference in time spent near the aversive cue <6 seconds per minute), but individual differences in learning ability did not explain substantial variability. We speculate that either the effect of aversive learning was too weak to quantify consistent individual differences, or directional selection might have eroded additive genetic variance. Finally, we discuss how confounded repeatability assays and publication bias could have inflated average estimates of repeatability in animal behaviour publications.

**Summary Statement:** Zebrafish exhibit low repeatability (intra-class correlation) in an aversive learning assay possibly due to past selection pressure exhausting genetic variance in this learning trait.

## Introduction

Animals use the cognitive process of learning, which can be defined as a change in behaviour due to past experience, to respond to the environment (Kawecki, 2010). Learning has a profound influence on survival and reproductive success (Krebs & Davies, 1987; Skinner, 1984), and has been studied in a wide range of taxa. For example, individual learning speed has been correlated with foraging performance in bees (Raine & Chittka, 2008) and grasshoppers (Pasquier & Grüter, 2016); and greater cognitive capacity has been linked to higher reproductive success in magpies (Ashton et al., 2018) and male robins (Shaw et al., 2019), as well as to healthier body condition in wild primates (Huebner, Fichtel, & Kappeler, 2018).

Animals learn through association, which is reinforced differently by positive and negative experiences (appetitive and aversive learning, respectively). Appetitive learning takes place when individuals associate a stimulus with a ‘positive’ event, usually a food reward stimulus, whereas in aversive learning the association is with a ‘negative’ event, usually a fear inducing stimulus. Failing to learn from positive experiences (appetitive learning) prevents a potential benefit (i.e. a minor opportunity cost). Failing to learn from negative experiences may yield an immediate fatal cost. Therefore, both types of learning can increase lifetime fitness and drive natural selection, but appetitive learning may be under weaker selection than aversive learning.

For traits to evolve they need heritable variation that can be subject to selection. For labile traits (i.e. traits expressed more than once over a lifetime) the consistency of individual differences in trait expression indicates potential heritability. The common approach to quantify consistent individual differences in eco-evolutionary studies is estimating the statistical index ‘repeatability’ *(R;* otherwise known as the ‘intra-class correlation coefficient’ or ICC; Lessells & Boag, 1987; Nakagawa & Schielzeth, 2010). Repeatability partitions variance into within-individual (residual) and between-individual components. Biologically, the repeatability of a trait is a measure showing the amount of observed variance which is due to individuals sustaining trait differences between each other (Nakagawa & Schielzeth, 2010), but estimates can be inflated by measurement errors and experimental confounds (Dohm, 2002; Niemelä & Dingemanse, 2017).

Generally, behavioural traits are moderately repeatable (*R* = 0.34; Bell et al., 2009; cf. Holtmann et al., 2017), with cognitive behavioural traits showing somewhat lower repeatability (*R* = 0.15 – 0.28; Cauchoix et al. 2018). Our understanding of how natural selection shapes the evolution of cognitive traits remains poor (Boogert et al., 2018). Despite the extensive literature on aversive learning, no published study has comprehensively quantified its repeatability (but note Cauchoix et al. (2018) includes three unpublished studies with some measures of aversive learning). To reduce this knowledge gap, we quantify the repeatability of aversive learning behaviour in zebrafish *(Danio rerio),* a popular model organism in cognitive science (Gerlai, 2016; Norton & Bally-Cuif, 2010). Zebrafish exhibit a range of distinct behaviours that can be measured in previously established assays (Fangmeier et al., 2018; Meshalkina et al., 2017).

Here, we use an avoidance conditioning assay — associating a visual cue with a mild electric shock — to thoroughly assess the repeatability of aversive learning in zebrafish. We expect individuals to consistently differ in their aversive learning speeds (i.e. separation of better and worse learners). First, we examine repeatability across different colour pairs (four different pairs with eight possible combinations: 8 measurements per individual). We expect individuals to show consistent differences in aversive learning ability and, given the estimates for appetitive learning summarised in Cauchoix et al. (2018), predict a low to moderate repeatability. Second, to examine whether a constant learning environment increases the consistency of individual differences, we examine repeatability within one colour pair (both combinations of green and blue; 3 repeated measurements per individual for each colour).

## Methods

### Zebrafish population

Adult wildtype zebrafish were bred and maintained at the Garvan Institute of Medical Research in Sydney, Australia. Fish were housed in 3.5L Tecniplast ZebTEC tanks (maximum of 24 fish per 3.5L tank) under standard laboratory conditions (~28°C; ~pH 7.5; ~1000 μs conductivity; 12/12h from 7:30 light/dark rotation) and fed live *Artemia salina* nauplii twice a day and commercially available fish food once per day (O.range GROW-L).

We marked juvenile fish for individual identification at around 90 days post-fertilisation with coloured tags (red, brown, purple, black, white, yellow, orange, pink, or green). For marking, fish were anesthetised in a tricaine solution (4.2ml of 0.4% in 100ml of system water) for 20 seconds before being injected with Visible Implant Elastomer tags (VIE, Northwest Marine Technologies, Inc.; Shaw Island, Washington, United States). We injected fish twice (unless one mark was blank), one on either side of the dorsal fin (Hohn & Petrie-Hanson, 2013). Among these marked fish, we used a total of 103 zebrafish with approximately equal sex ratios kept in 4 tanks of 24 individuals (12 males, 12 females) for both experiments. At any one time during the experiments, the same 96 fish were used, but to compensate for death, illness or experimenter error, seven fish were replaced by seven new fish over the three month study. Due to incomplete data for zebrafish size (described below) the across conditions and within conditions analyses included 93 and 94 zebrafish, respectively. The Garvan Animal Ethics Committee approved all procedures described above and experiments described below (ARA 18_18). Further, Garvan veterinarians oversaw fish welfare associated with aversive learning prior to our pilot tests.

### Experimental Design

#### Aversive Learning Assay

We used an avoidance conditioning method to quantify aversive learning in a simple, automated assay (Brock et al., 2017; Fontana et al., 2019). We ran all assays using four Zantiks AD units (Zantiks Ltd., Cambridge, UK; see Supplementary Figures 1 and 2). The units employed infrared tracking using an integrated computer to record fish movement and collect data. In the assay, a visual cue (colour or pattern) was associated with a negative stimulus (brief mild electric shock; 7V DC 80ms), which motivated fish to avoid the associated visual cue. We then measured the extent of avoidance (i.e. time spent away from the cue associated with an electric shock) compared to the baseline preference to quantify aversive learning (association response). We based our initial assay parameters (e.g., the acclimation period, voltage, etc) on previous research (Brock et al., 2017), and subsequently modified the parameters based on the outcomes of pilot tests (see Supplementary Information).

Before each assay we individually placed fish into one of four lanes within rectangular tanks (see Figure 1A). For the assay, we exposed the fish to four stages; (i) Acclimation: we habituated the fish to isolation in a novel environment over a 30-minute acclimation period (Figure 1B); (ii) Baseline: the tank was visually split into two even zones via the colour displaying screen at the bottom of the tank (Figure 1C). One of these two colours would later become conditioned with the mild electric shock (CS+), the other colour remained unconditioned (CS-). Here, the position of the colours (left or right) automatically switched every five minutes for a period of 30-minutes and we recorded zebrafish preference for the CS+ to obtain a baseline preference before conditioning; (iii) Conditioning: first, the CS+ (visual cue associated with shock) was displayed across the entire screen for 1.5 seconds then immediately afterwards paired with the US (mild electric shock) to condition the fish to an aversive experience. Second, the CS- (visual cue not associated with shock) covered the screen for 8.5 seconds (Figure 1D). This phase was repeated nine times, sufficient for fish learning to avoid the CS+; and, (iv) Probe: akin to the baseline period, the tank was split into two even zones (left or right) depicted by different visual cues. We tracked fish movement and recorded fish preference for the visual cue associated with the shock (CS+) over 5 minutes. During this time, the visual cues switched every minute (see Figure 1E). Probe CS+ preference was used in comparison to baseline CS+ preferences to quantify learning. We used only 2 minutes out of the 5-minute probe time since we determined in our observations (see Supplementary Figure 5 & 6) a clear decrease in learning response. This probe length is similar to other studies Brock et al. (2017) use a 2-minute probe and Fontana et al., (2019) use a 1 minute probe.

**Figure 1.**
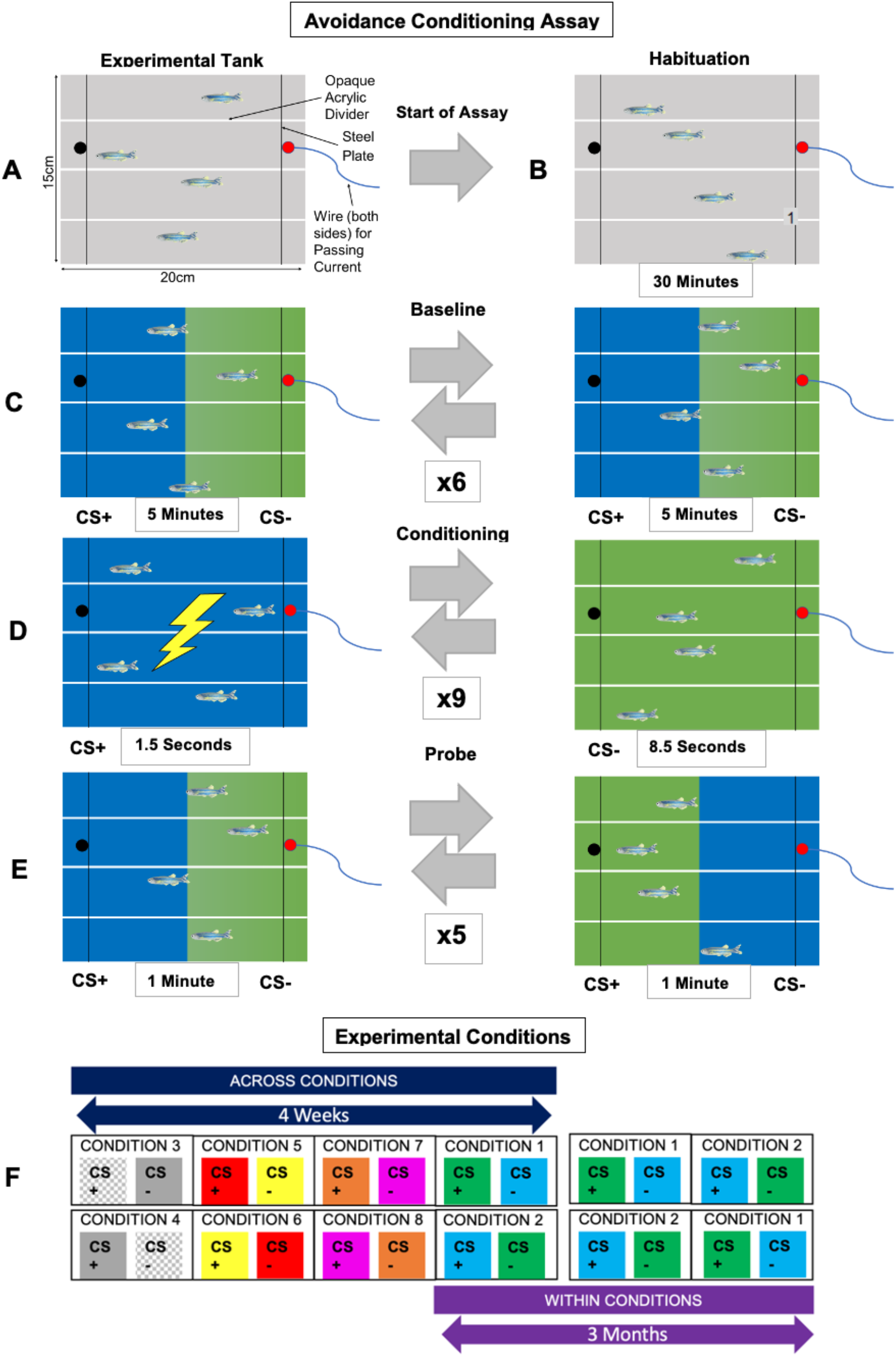
Colour conditions and aversive learning assay: (**A**) zebrafish are placed in the experimental tanks and (**B**) acclimated to the novel environment for 30-minutes; (**C**) in a 30-minute baseline period, initial CS± preference is established; (**D**) during the conditioning phase, fish are presented the CS+, then immediately subjected to a mild electric shock; and (**E**) in a 5-minute probe phase, learning is determined by fish spending less time in the CS+ when compared to the baseline. (**F**) Each condition is a combination of two visual cues (zones), one conditioned to a mild electric shock (CS+), the other is not (CS-). Across conditions eight colour conditions and eight sessions (each session is represented by a white box). Within conditions: two colour conditions and four sessions (in addition to two sessions in Experiment 1).

#### Experimental Conditions

We used a range of colour conditions to test aversive learning. Each condition was comprised of two visual cues, one aversive and one control (CS+ paired with CS-) (Figure 1F). We selected different colour combinations to use as visual cues for the zebrafish, which had either been worked in pre-existing assays or were reported to evoke a clear colour preference (Brock et al., 2017; Roy et al., 2019). As a result, we chose seven colours (green, blue, grey, orange, magenta, red, yellow) and 1 pattern (check; hereafter, this pattern is also referred to as a ‘colour’ with the others). We used four visual cue combinations (‘Check/Grey’, ‘Green/Blue’, ‘Red/Yellow’, ‘Magenta/Orange’) and their reverse (‘Grey/Check’, ‘Blue/Green’, ‘Yellow/Red’, ‘Orange/Magenta’) for a total of eight conditions. For example, the ‘Check/Grey’ condition used check pattern as the CS+ (cue associated with shock) and grey colour as the CS- (control cue); the ‘Grey/Check’ condition used grey colour as the CS+ and check pattern as the CS-, and so on.

Prior to the experiment, we assigned fish into quartets (four fish that underwent trials within the same Zantiks unit/assay tank simultaneously) that systematically rotated between trials. The balanced design accounted for three potential confounding variables: the time of day (quartet rotated), Zantiks unit (quartet rotated), and lane position (individual within quartet rotated). We estimated repeatability in two different situations (across conditions and within a single condition). Across conditions, we ensured fish experienced trials from all four colour pairs before subjecting them to their exact reverse four conditions (with trials conducted over four weeks in June and July 2019). We included this form of reverse learning to negate memory of the CS+ colour between trials, which may impact both baseline and probe colour preference. Within conditions, each zebrafish underwent trials in the ‘Blue/Green’ and ‘Green/Blue’ conditions a further two times (over two weeks in September 2019).

#### Fish Size Measuring

We took photos of each fish approximately one week after across-conditions trials and another set of photos approximately one week after within-conditions trials. We captured top down photos of live fish and measured fish in ImageJ (Schindelin et al., 2015). We used fish length (standard length) and width (at widest part of body) to calculate the ellipsoid size of the fish by using 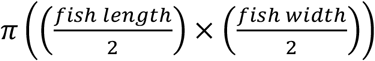. This controlled for a potential size effect resulting from loss of penetrance and effectiveness of the mild electric shock due to larger body size.

### Data Processing and Analysis

All data processing and analyses were conducted in the *R* computing environment (version 4.0.2; R Core Team, 2019). Linear mixed models were run using the *lme4* package (version 1.1.21; Bates et al., 2014) in conjunction with the lmerTest package (version 3.1.2; Kuznetsova, Brockhoff, & Christensen, 2017), that provides Satterthwaite’s degrees of freedom correction. We obtained repeatability values via the *rptR* package (version 0.9.22; Stoffel et al., 2017) that uses the *lme4* pacakge to run mixed models. Based on visual assessments of residual distributions, assumptions of normality and constant variance were not clearly violated. The Zantiks units recorded time spent in each CS zone, total distance travelled and how often fish changed zones. All code, and the raw and processed data, are available at: https://osf.io/t95v3/. We deemed our results statistically significant at the alpha = 0.05 level (or when 95% confidence intervals did not overlap zero).

#### Quantifying Aversive Learning

We determined learning by the difference in time that fish spent in the CS+ before and after the aversive experience. To analyse learning across all the sessions included in this study, we used the time difference *(‘difference’ = time spent in the CS+ during baseline – time spent in the CS+ during probe)* as the response variable in a linear mixed-effects model (LMM) via the *lmer* function in the *lme4* package. We fitted individual ‘fish ID’ 96 levels) and ‘experimental condition ID’ (8 levels, see Figure 1F) as random effects in the model. Also, we included the following fixed effects: (1) ‘sex’ (female or male) to investigate sex differences in learning, (2) ‘day’ since first trial, to account for time effects of sequential days on learning or learning via repeated trials (e.g., 1 being the first day and 8 being the 7^th^ day from the first), (3) ‘fish size’ to control for fish’s response to conditioning which might be size dependent due to potential differences in body penetrance of a mild shock, (4) ‘learning’ (initial and reverse) to find if learning was affected when the CS± of a condition were switched in successive trials. Note that we *z*-transformed the fixed effects ‘day’ and ‘fish size’ to make the intercept meaningful and slope estimates comparable (Schielzeth, 2010).

#### Quantifying the Repeatability of Aversive Learning

We obtained enhanced agreement repeatability (hereafter referred to as repeatability) estimates by incorporating statistically significant fixed effects from the model and retaining their variance in the denominator (Nakagawa & Schielzeth, 2010). We only fitted the random effect ‘fish ID’ and included ‘sex’ as a fixed effect. The R package *rptR* computes repeatability values using the within and between individual variance in linear mixed models fitted with restricted maximum likelihoods (Nakagawa & Schielzeth, 2010). Using *rptR,* we obtained standard errors and 95% confidence intervals (CIs), each model set to 10,000 bootstrap samples. Following Bell (2009) and Wolak (2012), we categorised our repeatability results into low (<0.2), moderate (>0.2 – <0.4) and high (>0.4).

#### Colour Preference and Repeatability

An underlying assumption of our aversive learning assay was that zebrafish have the ability to discriminate between different colours. Therefore, from the baseline period (prior to aversive conditioning), we quantified underlying colour preferences (tendency to associate more heavily with one colour in a pair), and the consistency of individual differences in colour preference (i.e. repeatability of colour preference).

In each condition, preference for one colour was only compared to the other paired colour (e.g. preference for red is only relative to preference for yellow; see Figure 1F). Given we examined relative colour preference, preferences for either colour in a condition were the inverses of each other. Hence, to be able to determine colour preference for each colour, we grouped conditions of matching colours into four groups for analysis (e.g. Group 1, ‘Red/Yellow’ & ‘Yellow/Red’; Group 2, ‘Green/Blue’ & ‘Blue/Green’; Group 3, ‘Check/Grey’ & ‘Grey/Check’; Group 4, ‘Orange/Magenta’ & ‘Magenta/Orange’).

To analyse relative colour preference, we ran LMMs for each group of colours using across conditions data. We used baseline colour preference as the response variable ‘baseline’ for these models. We fitted the random effect ‘fish ID’ in the models (Group 1 & 4, 97 levels; Group 2 & 3, 98 levels; levels differ because one fish died prior to completing all conditions). Further, we fitted the following fixed effects: (1) ‘day’ (days since first trial) to control for potential colour preference change with time, (2) ‘sex’ (male or female) to account for sex differences and (3) ‘learning’ (initial and reverse) to see the effect of reverse learning on colour preference. To determine the repeatability of colour preference, we used *rptR* mixed-effects models with the response variable ‘baseline’ to generate repeatability estimates. We did not find any fixed effects to be statistically significant, as such, they were excluded, and the colour preference models were fit with the random effect ‘fish ID’.

## Results

### Do Zebrafish Show Appropriate Responses in an Aversive Learning Assay?

Zebrafish spent more time avoiding the CS+ following conditioning, showing evidence of learning (across conditions: female average = 3.89 seconds per min, SE = 1.05, t_33_ = 3.65, *P* < 0.001; male average = 5.64 seconds per min, SE = 0.94, t_22_ = 5.21, *P* < 0.001; Figure 2B). Overall, males avoided the CS+ more than females, but this result was not statistically significant (1.75 seconds per min, SE = 0.90, t_108_ = 1.93, *P* = 0.055). Reverse learning had a non-significant slight negative effect (−1.11 seconds per min, SE = 1.03, t_1008_ = −1.07, *P* = 0.281). All other fixed effects did not significantly impact learning (see Supplementary Table 3 for all model outputs).

**Figure 2.**
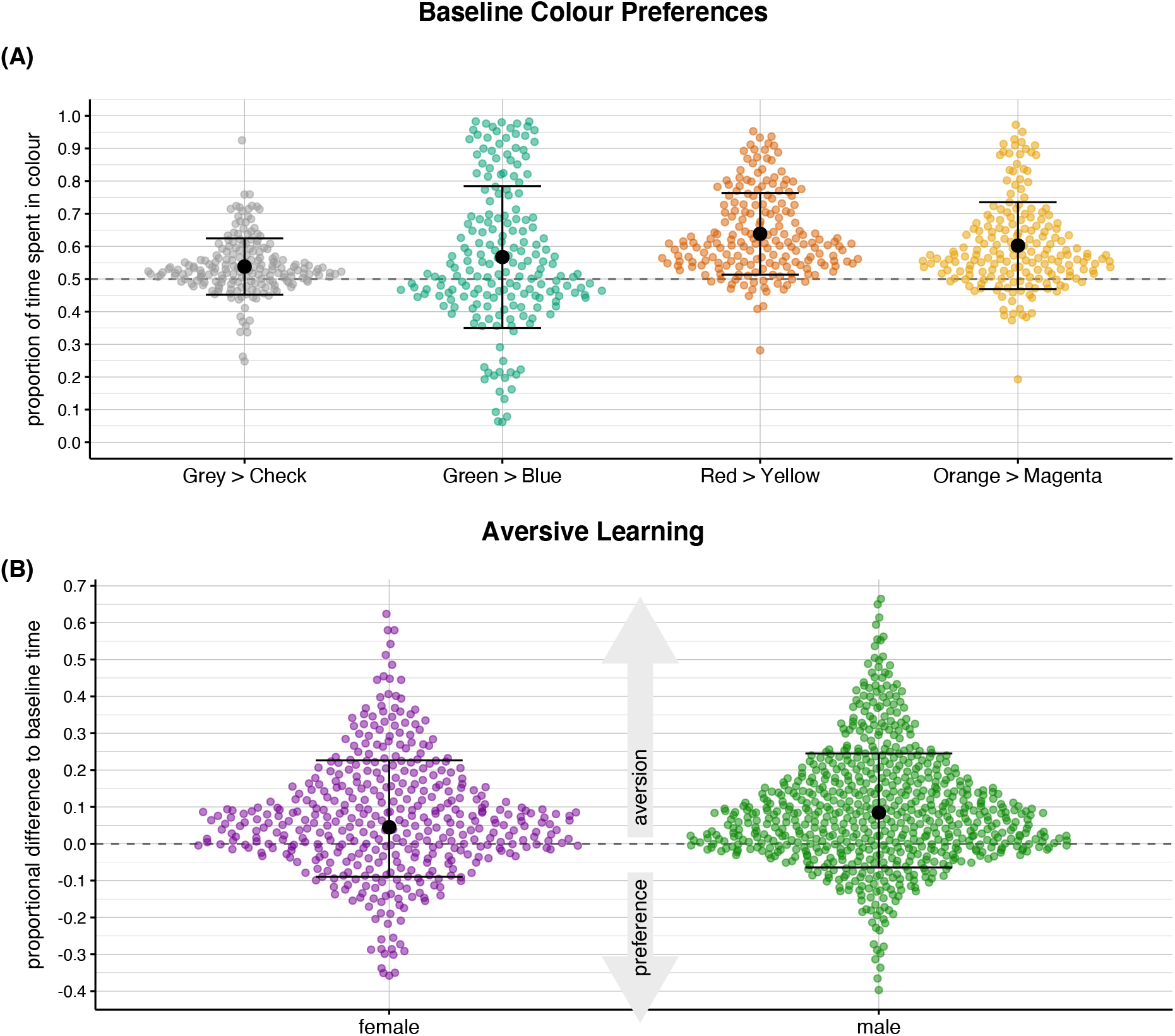
Violin plots for colour preferences and aversive learning. Smaller coloured points depict individual trials. Larger black points and error bars depict means and standard deviations of observations. (**A**) The top panel shows the tendency of zebrafish to favour one colour in a pair during the baseline period (i.e. before administration of electric shocks). The dashed horizontal line at 0.5 represents no colour preference (i.e. spending 30 seconds in each colour zone). (**B**) The bottom panel shows means and variation in aversive learning, split by sex (female = purple; male = green) when all the session data is combined. Points above the line at zero depict trials in which zebrafish spent less time in the aversive stimulus colour in the probe period (the colour associated with an electric shock) relative to the baseline period (i.e. aversive learning).

### Is Aversive Learning Repeatable Across and Within Conditions?

We found very low repeatability across the eight different conditions (*R* = 0.04, 95% CI [0.001 – 0.097], Figure 3B). Within conditions, the repeatability (point-estimate) of the ‘Green/Blue’ condition was even lower than the across-condition estimate (*R* = 0.02, 95% CI [0 – 0.153]), while repeatability was higher in the ‘Blue/Green’ (*R* = 0.15, 95% CI [0.023 – 0.278]; see also Supplementary Figure 3 for male and female estimates).

**Figure 3.**
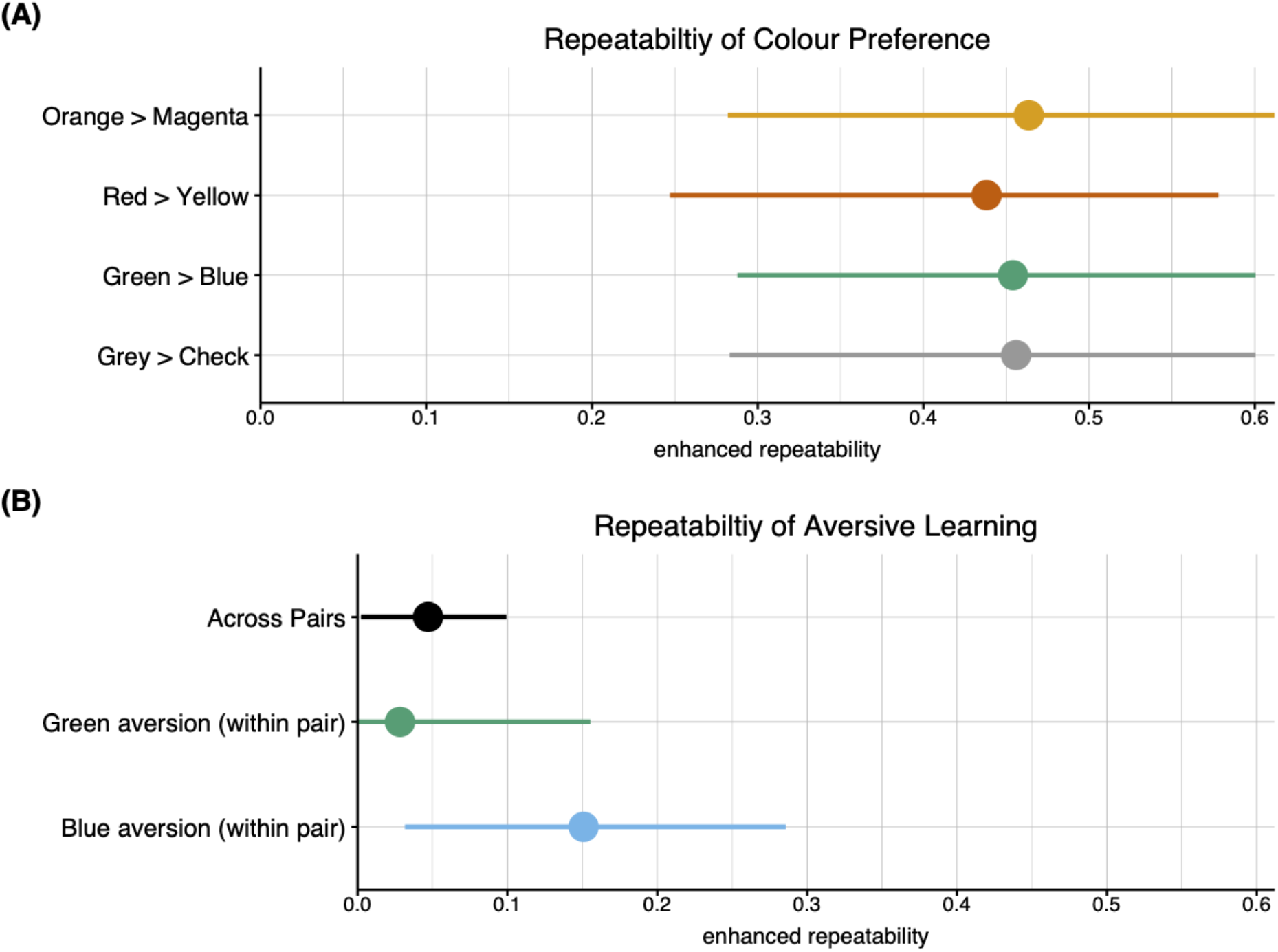
Repeatability of colour preference and aversive learning in zebrafish. Points and whiskers represent means and 95% confidence intervals via parametric bootstrapping. (**A**) Zebrafish show consistent individual differences in colour preferences (variation depicted in Figure 2). (**B**) Zebrafish show somewhat consistent individual differences in aversive learning within the Blue/Green pair, but not within the Green/Blue pair or across all colour combinations.

### Do Zebrafish Display Colour Preferences and is Preference Repeatable?

Zebrafish showed strong relative colour preference in all four conditions (see Figure 2B). In addition, fish exhibited repeatable relative colour preferences which were highly consistent across all four conditions (Figure 3A; Grey: *R* = 0.45, 95% CI [0.276 – 0.607]; Green: *R* = 0.45, 95% CI [0.278 – 0.604]; Red: *R* = 0.43, 95% CI [0.250 – 0.584]; Orange: *R* = 0.46; 95% CI [0.283 – 0.605]; see Supplementary Table 1 and 2 for all repeatability estimates).

## Discussion

We investigated aversive learning in zebrafish and quantified repeatability in two scenarios. We first tested if fish displayed stable individual differences across different learning environments, equivalent to methods investigating ‘animal personality’ (i.e. consistent differences over time and contexts; Sih et al., 2004). We found negligible repeatability in aversive learning across conditions, despite individuals being able to discriminate between colours (as measured by a moderate repeatability of colour preferences). Then, we examined repeatability within two separate conditions, which is more consistent with the idea of ‘pseudo-repeatability’ (where consistency is inflated due to measurements under an identical condition; Niemelä & Dingemanse, 2017). Within two conditions, we found negligible repeatability in one condition (‘Green/Blue’ *R* = 0.02), and low repeatability in the other (‘Blue/Green’ *R* = 0.15; Figure 3B). Therefore, the substantial variation in aversive learning we observed was most likely driven by current (intrinsic or extrinsic) environmental factors, rather than additive genetic variance or canalized developmental differences (cf. Sznajder, Sabelis, & Egas, 2012)

Our results are surprising, given low to moderate repeatability of behaviour and cognition reported in two meta-analyses. For behaviour generally, Bell et al (2009) reported an average repeatability of *R* = 0.34. For cognitive performance, Cauchoix et al. (2018) found *R* = 0.15-0.28, mostly based on temporal repeatability estimates from appetitive learning trials. Below we discuss four potential reasons why zebrafish in our experiment showed much less consistent individual differences in average learning compared to those previous estimates from Cauchoix et al. (2018) and Bell et al. (2009).

First, while zebrafish did demonstrate aversive learning, the effect was small, and in many trials, individuals did not seem to avoid the negative stimulus, perhaps due to not learning or quickly forgetting; on average, individuals spent 3.89 (females) and 5.64 (males) fewer seconds per minute respectively in the negatively associated colour following conditioning (Figure 2B). Low repeatability could therefore be caused by zebrafish being largely insensitive to the conditioning (i.e. bad aversive learners, or a weak assay). However, the fact that there was a population shift in the direction of aversive learning raises the question of why individuals who learnt in one trial did not maintain their performance across trials; if a particular subset of zebrafish had consistently learnt, or failed to learn, then we would have detected higher repeatability. Further, while the behaviour change following aversive conditioning was modest, zebrafish learnt much faster (in 1.5 minutes) compared to previous assays with appetitive training (e.g., over 20 days; Brocks et al. 2017). As far as we are aware, no studies have investigated a relationship between the strength of associative learning and the magnitude of repeatability.

Second, past selection pressures on our study population may have eroded additive genetic variance associated with aversive learning, which was not restored in the intervening generations. In the wild, aversive learning could be under strong selection (e.g. to learn to evade predators), and individuals could be selected to learn from negative experiences as quickly as possible. Indeed, aversive learning could be under stronger selection than appetitive learning, as mortality costs of negative experiences can easily exceed opportunity costs of missing positive experiences. Stronger selective pressures could explain why we found substantially lower repeatability for aversive learning compared with previous results for appetitive learning. In a similar vein, a trait more closely associated with fitness (e.g., aversive learning) tends to not be as heritable (thus, repeatable; cf. Dohm, 2002) than less fitness related traits (e.g., appetitive learning; Merilä & Sheldon, 2000). However, we cannot be sure that whether the performance of zebrafish in our laboratory assay accurately captures their ability to aversively learn in their natural habitat.

Third, some of the repeatability values in the meta-analyses by Cauchoix et al. (2018) and Bell et al. (2009) may have been overestimated. An inflated repeatability estimate, also known as ‘pseudo-repeatability’, is the result of within-individual variation being erroneously accredited to differences between individuals (Niemelä & Dingemanse, 2017; Westneat, Hatch, Wetzel, & Ensminger, 2011). Pseudo-repeatability occurs when the conditions between measurements are too similar (e.g., environmental conditions are unchanged or intervals between measurements are too short), and might explain why we found higher repeatability when zebrafish were measured repeatedly within a single condition (‘Blue/Green’; *R* = 0.15), compared to across eight separate conditions (although no inflation was seen in ‘Green/Blue’). On closer inspection, some of studies in Cauchoix et al. (2018) and Bell et al. (2009) included testing conditions which did not change over the course of a study, similar to our within-condition estimates. Further, both Cauchoix (2018) predominately included studies with intervals under a week and Bell et al. (2009) almost all were under a year. Bell et al. (2009) reported that short intervals between measurements were signficantly associated with higher repeataibltiy values in line with pseudo-repeatability. Relevantly, two recent studies on birdsong reported that associative learning among individuals was not repeatable between years, indicating that estimates obtained over short intervals may not be a true reflection of phenotypic constancy defined in animal personality (Soha et al., 2019; Zsebők et al., 2017).

Fourth, the meta-analytic repeatability estimates by Bell et al. (2009) might have been overestimated due to a potentially widespread publication bias in the literature reporting repeatability of behaviour (cf. Parker et al., 2016). Our across conditions repeatability estimate is markedly low in comparison to that of general behaviour founded in Bell et al. (2009; *R* = 0.34) that only included published studies. Cauchoix et al. (2018) included many unpublished datasets (n = 38) compared to published datasets (n = 6); they mentioned that their unpublished datasets produced, overall, a lower repeatability estimate than that of the published studies. This finding is consistent with the pattern that larger effect sizes are more likely to be published. It is possible that publication bias has further contributed to an inflation of the overall repeatability estimates in the published literature. However, recent studies are increasingly reporting non-significant and low repeatability (e.g., Reichert et al., 2020; Vernouillet & Kelly, 2020). Therefore, an updated future meta-analysis may reveal a lower overall repeatability estimate in behaviour.

In conclusion, zebrafish did not show clear consistent between-individual differences in aversive learning. The low repeatability could potentially indicate that strong past selection pressure has almost driven aversive learning to fixation, because of the vital importance to learn to avoid danger. In addition, many researchers may have unknowingly included confounded pseudo-repeatability results in their studies. In turn, inflating published repeatability estimates and presenting the repeatability of behaviour and learning-associated behaviour higher than the ‘true’ repeatability of behaviour. Further, a bias to withhold non-significant findings from publishing may have exacerbated this inflation in the literature. We contend that these issues can be diminished in future behavioural research by controlling for confounding effects and reporting every estimate of behavioural traits, whether repeatable or not.

## Acknowledgement

We are grateful for the staff at the Biological Testing Facility, Garvan Institute of Medical Research for their support and husbandry of zebrafish. This study was supported by ARC (Australian Research Council) Discovery grant (DP180100818). No competing interests declared.

## Supplementary Information (SI)

### Supplementary Methods

#### Zantiks Experimental Units

We used Zantiks AD fully automated units to conduct our behavioural experiments (Zantiks Ltd., Cambridge, UK; Supplementary Figure 1). The design enabled comprehensive standardised cognitive assays on zebrafish. The boxes’ capabilities include infrared tracking, a stimulus screen, feeding mechanisms, removable tanks with modifiable inserts, an in-built computer, console interface and video recording. They were well equipped to conduct simple experimental manipulation and provide a range of stimuli (colours, patterns or images) to measure behavioural responses.

During experiments, we placed portable tanks (length 20cm: height 14cm: width 15cm; 2.6L system water; see Supplementary Figure 2, picture C) containing the fish onto the screens inside the units (see Supplementary Figure 2, picture B). We presented experimental stimuli via the screen through the transparent base of the tank. Fish location co-ordinates were tracked via an inbuilt infrared (IR) camera situated at the ceiling of the unit and another IR source underneath the screen. A basal screen enabled a near completely closed system inhibiting external disturbances.

#### Pilot Experiments

To find the best parameters to use in the avoidance task, we carried out numerous pilot assays. Specifically, we examined three parameters: stimulus type (colour or pattern), assay length and voltage. Our aim was to find the shortest assay length and lowest voltage suitable to yield a behavioural response from the fish. At the same time we determined which stimuli (colours) would be ideal, testing stimuli used in the literature (Brock et al., 2017) and two colours that we did not find in the literature, orange and magenta.

With guidance from previous research (Brock et al., 2017), we conducted preliminary tests to identify the lowest voltage setting required to yield an adequate learning response. We tested three settings: five, seven and nine volts, each applied nine times per trial. The seven-volt setting elicited the most fish avoidance for the CS+ in the probe stage (see Supplementary Figure 4). Based on this finding we used seven volts applied nine times as the US in the conditioning phase for our experiments.

The previously developed assay by Brock et al. (2017) comprised of 3 stages: baseline, conditioning and probe. We extensively tested the three stages to decide the optimal length for each. Akin to other studies, the baseline and conditioning stages remained the same length (Brock et al., 2017; Fontana et al., 2019). However, we lengthened the probe period from two to five minutes to provide a wider range for potential analysis. Further, we introduced an acclimation stage to ensure a consistent association response from the fish (Thomson et al., 2020). The acclimation stage is absent in some studies, although when present can range in length from 10-minutes (Baker & Wong, 2019; Kenney et al., 2017) to over the course of multiple days (Kaneko, Masuda, & Yamashita, 2019; Namekawa, Moenig, & Friedrich, 2018). Following the data collected during our pilot assays, we found a 30-minute acclimation period just prior to data collection afforded the best association response.

Our pilot results indicated a steep decline in association response after two minutes in the probe period. Although these results aligned with the literature (Brock et al., 2017 2-minute probe; Fontana et al., 2019 1-minute probe) we integrated an extended probe period (the probe phase above) to verify if fish would display a similar deterioration. As expected, the fish exhibited a corresponding drop in association response after two minutes. Consequently, for our analysis we confined the extended probe period to two minutes since the ensuing deterioration may indicate memory loss or habituation to the CS+ post conditioning or a new learning event.

### Supplementary Notes

#### Sex Differences in Repeatability

We found males we more generally more repeatable than females (Figure S3) in aversive learning. We found this result across conditions (males, sample size = 63, *R* = 0.06, 95% CI [0.007 – 0.091]; females, sample size = 46, *R* = 0.00, 95% CI [0 – 0.055]) and in the ‘Blue/Green’ condition (males, sample size = 62, *R* = 0.23, 95% CI [0.050 – 0.374]; females, sample size = 37, *R* = 0.02, 95% CI [0 – 0.195]). This result was not anticipated since females are reported to be more repeatable than males in behaviour (Bell et al., 2009). We found no statistically significant difference in repeatability between males and females, displayed by no overlap over zero in bootstrap distribution displayed in Supplementary Figure 4.

In relative colour preference, we found males were more repeatable than female in the colours red, grey and orange but not green. Since we are the first to assess the repeatability of colour preference in zebrafish, we cannot compare to the literature, however, the sex differences in colour preference repeatability are mostly consistent with those in the repeatability of aversive learning.

### Supplementary Tables

**Supplementary Table 1.** Repeatability values for different conditions with bootstrapped 95% confidence intervals. All conditions display sexes mixed at the top then male and female results. Estimates with CIs that do not overlap zero are presented in bold.

| Conditions | Repeatability (*R*) | 95% Confidence Interval |
| --- | --- | --- |
| Across | 0.047 | 0.007 - 0.091 |
| Across Male | 0.069 | 0.014 - 0.154 |
| Across Female | 0 | 0 - 0.055 |
| ----- |  |  |
| Green/Blue | 0.028 | 0 - 0.137 |
| Green/Blue Male | 0.039 | 0 - 0.222 |
| Green/Blue Female | 0.016 | 0 - 0.203 |
| ----- |  |  |
| Blue/Green | 0.150 | 0.023 - 0.308 |
| Blue/Green Male | 0.232 | 0.050 - 0.374 |
| Blue/Green Female | 0.022 | 0 - 0.195 |
| ----- |  |  |
| First | 0 | 0 - 0.189 |
| First Male | 0 | 0 - 0.227 |
| First Female | 0.029 | 0 - 0.383 |
| ----- |  |  |
| Second | 0.012 | 0 - 0.200 |
| Second Male | 0 | 0 - 0.000 |
| Second Female | 0.073 | 0 - 0.419 |
| ----- |  |  |
| Third | 0 | 0 - 0.222 |
| Third Male | 0 | 0 - 0.275 |
| Third Female | 0 | 0 - 0.355 |
| All | 0.013 | 0 - 0.072 |
| All Male | 0.046 | 0 - 0.141 |
| All Female | 0 | 0 - 0.100 |

**Supplementary Table 2.** Repeatability estimates of relative colour preference with bootstrapped 95% CIs for red, green, grey and orange. Male and female preference included. Estimates with CIs that do not overlap zero are presented in bold.

| Colour | Repeatability (*R*) | 95% Confidence Interval |
| --- | --- | --- |
| Red | 0.438 | 0.250 - 0.584 |
| Red Male | 0.492 | 0.288 - 0.656 |
| Red Female | 0.331 | 0.009 - 0.586 |
| Green | 0.454 | 0.278 - 0.604 |
| Green Male | 0.434 | 0.215 - 0.614 |
| Green Female | 0.490 | 0.203 - 0.702 |
| Grey | 0.455 | 0.276 - 0.607 |
| Grey Male | 0.499 | 0.309 - 0.657 |
| Grey Female | 0.391 | 0.056 - 0.635 |
| Orange | 0.463 | 0.283 - 0.605 |
| Orange Male | 0.519 | 0.280 - 0.681 |
| Orange Female | 0.411 | 0.083 - 0.649 |

**Supplementary Table 3.**
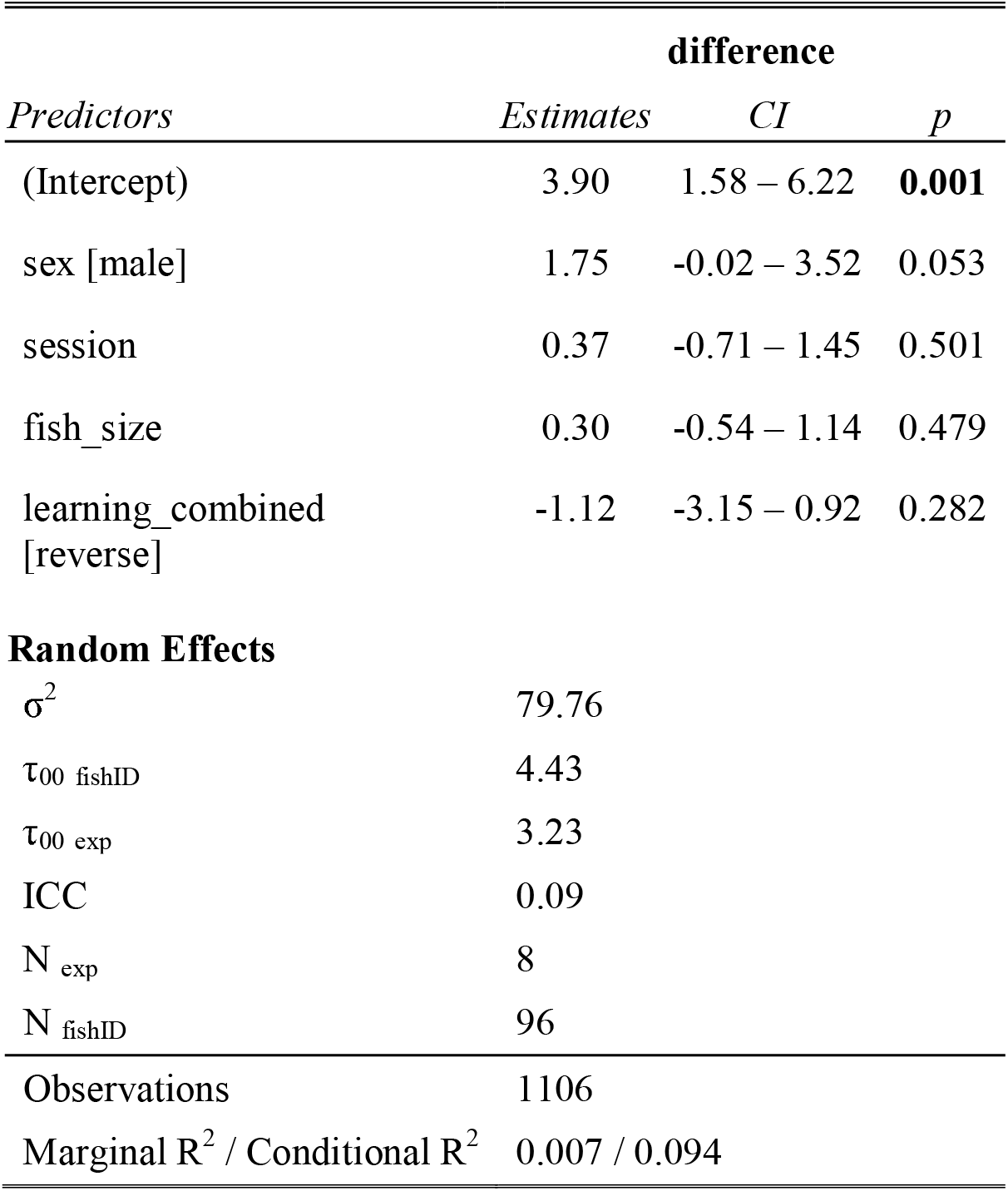
The outputs of fixed and random effects from the across conditions aversive learning mixed effect model. Significant results are displayed in bold.

| *Predictors* | **difference** |  |  |
| --- | --- | --- | --- |
|  | *Estimates* | *CI* | *p* |
| (Intercept) | 3.90 | 1.58 – 6.22 | **0.001** |
| sex [male] | 1.75 | -0.02 – 3.52 | 0.053 |
| session | 0.37 | -0.71 – 1.45 | 0.501 |
| fish_size | 0.30 | -0.54 – 1.14 | 0.479 |
| learning_combined<br>[reverse] | -1.12 | -3.15 – 0.92 | 0.282 |
| **Random Effects** |  |  |  |
| $\sigma^2$ | 79.76 | | |
| $\tau_{00}$ fishID | 4.43 | | |
| $\tau_{00}$ exp | 3.23 | | |
| ICC | 0.09 |  |  |
| $N_{exp}$ | 8 | | |
| $N_{fishID}$ | 96 | | |
| Observations | 1106 |  |  |
| Marginal $R^2$ / Conditional $R^2$ | 0.007 / 0.094 | | |

**Supplementary Table 4.** The outputs of fixed and random effects from the across conditions red colour preference mixed effect model. Significant results are displayed in bold.

| BASELINE |  |  |  |
| --- | --- | --- | --- |
| Predictors | Estimates CI |  | p |
| (Intercept) | 36.35 | 33.03 – 39.67 | <0.001 |
| day | -1.05 | -3.80 – 1.71 | 0.457 |
| sex [male] | 1.51 | -1.13 – 4.15 | 0.262 |
| learning [reverse] | 2.05 | -3.09 – 7.18 | 0.435 |
| Random Effects |  |  |  |
| $\sigma^2$ | 32.10 | | |
| $\tau_{00 \text{ fishID}}$ | 24.72 | | |
| ICC | 0.44 |  |  |
| $N_{\text{fishID}}$ | 98 | | |
| Observations | 192 |  |  |
| Marginal $R^2$ / Conditional $R^2$ | 0.014 / 0.443 | | |

**Supplementary Table 5.**
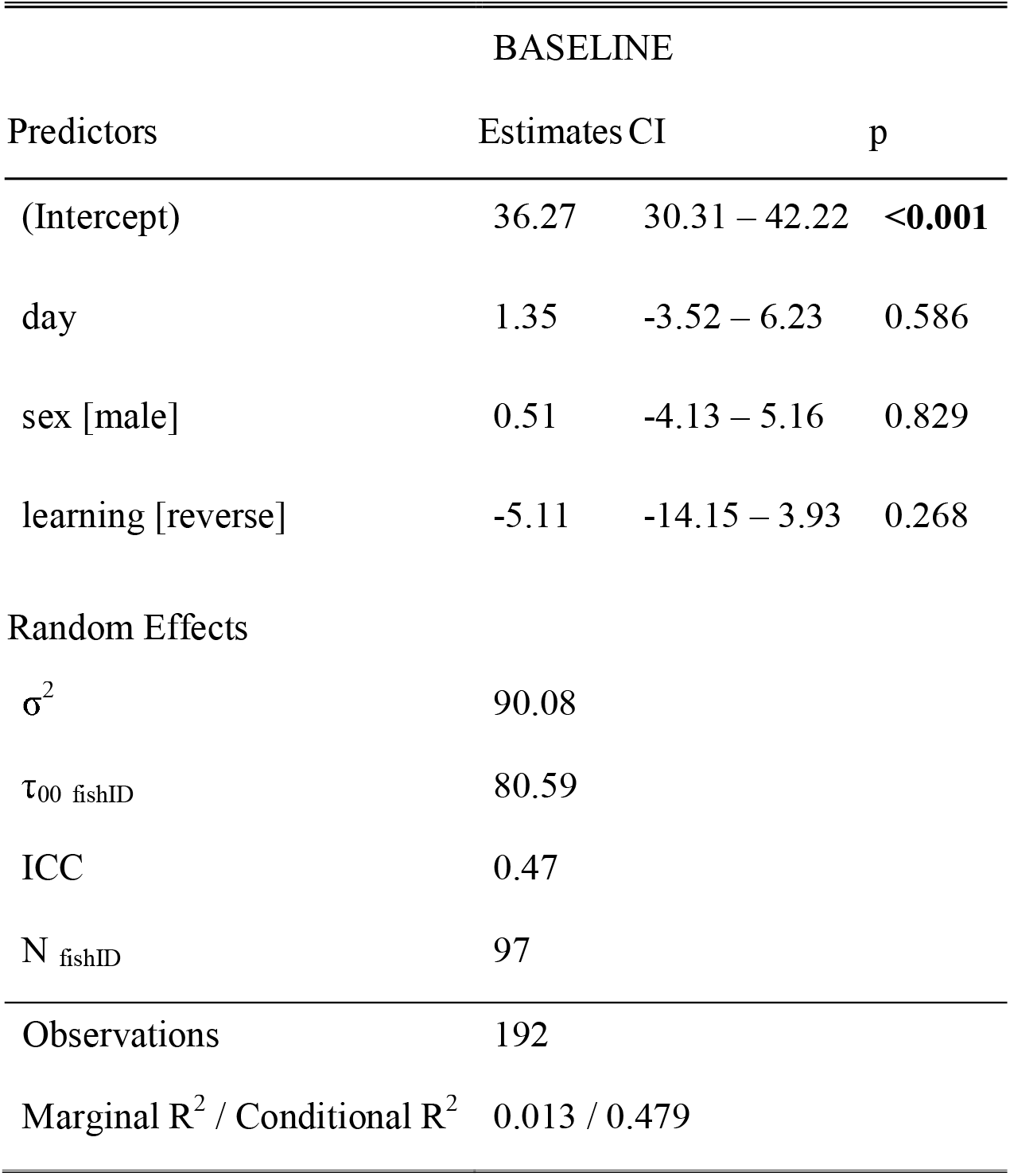
The outputs of fixed and random effects from the across conditions green colour preference mixed effect model. Significant results are displayed in bold.

**Supplementary Table 6.** The outputs of fixed and random effects from the across conditions check colour preference mixed effect model. Significant results are displayed in bold.

| BASELINE |  |  |  |
| --- | --- | --- | --- |
| Predictors | Estimates CI |  | p |
| (Intercept) | 32.81 | 30.54 – 35.07 | **<0.001** |
| day | 0.42 | -1.51 – 2.36 | 0.667 |
| sex [male] | -0.63 | -2.47 – 1.21 | 0.503 |
| learning [reverse] | -0.26 | -3.85 – 3.34 | 0.889 |
| Random Effects |  |  |  |
| $\sigma^2$ | 14.61 | | |
| $\tau_{00 \text{ fishID}}$ | 12.47 | | |
| ICC | 0.46 |  |  |
| $N_{\text{fishID}}$ | 97 | | |
| Observations | 192 |  |  |
| Marginal $R^2$ / Conditional $R^2$ | 0.007 / 0.464 | | |

**Supplementary Table 7.**
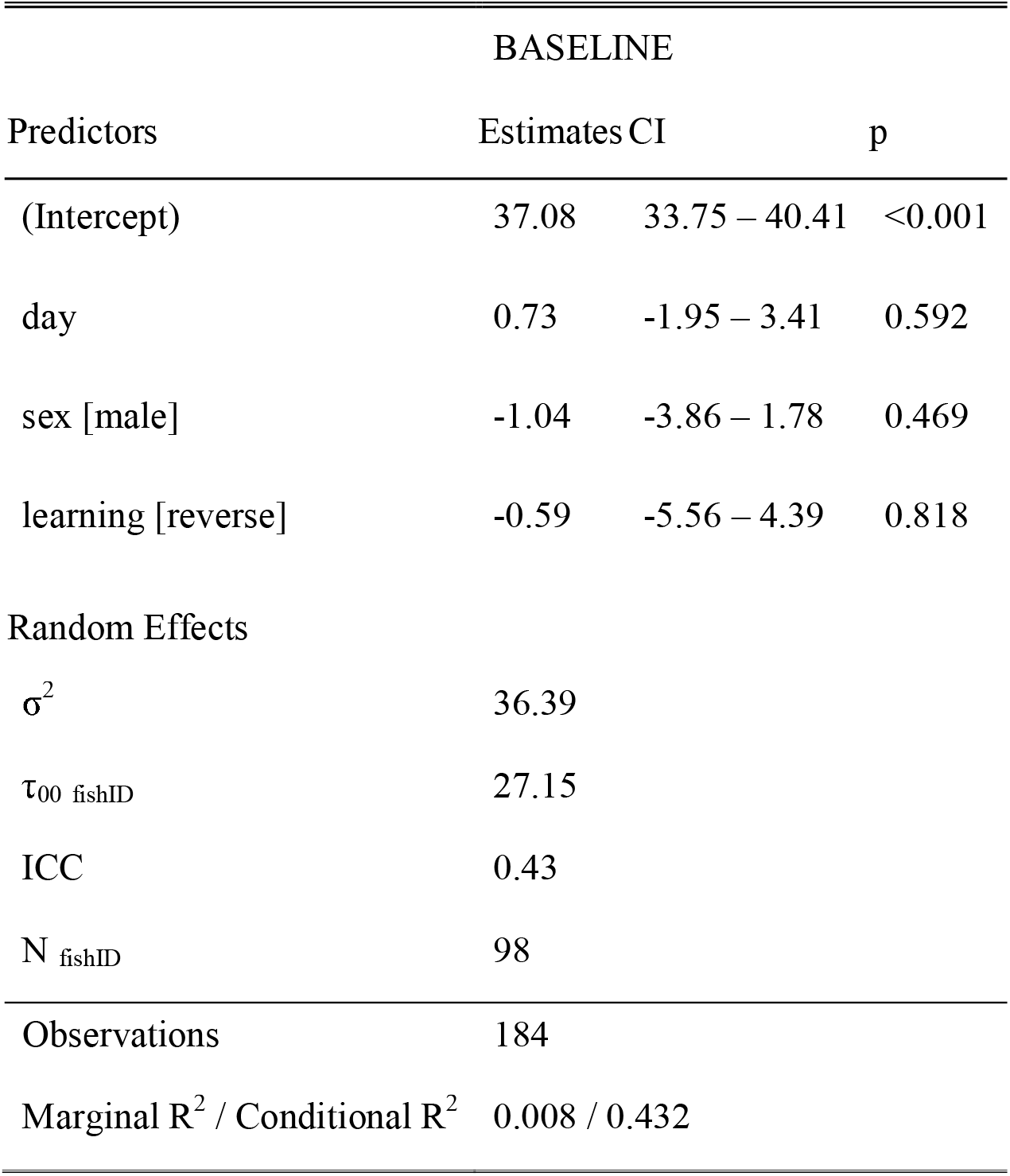
The outputs of fixed and random effects from the across conditions orange colour preference mixed effect model. Significant results are displayed in bold.

### Supplementary Figures

**Supplementary Figure 1.**
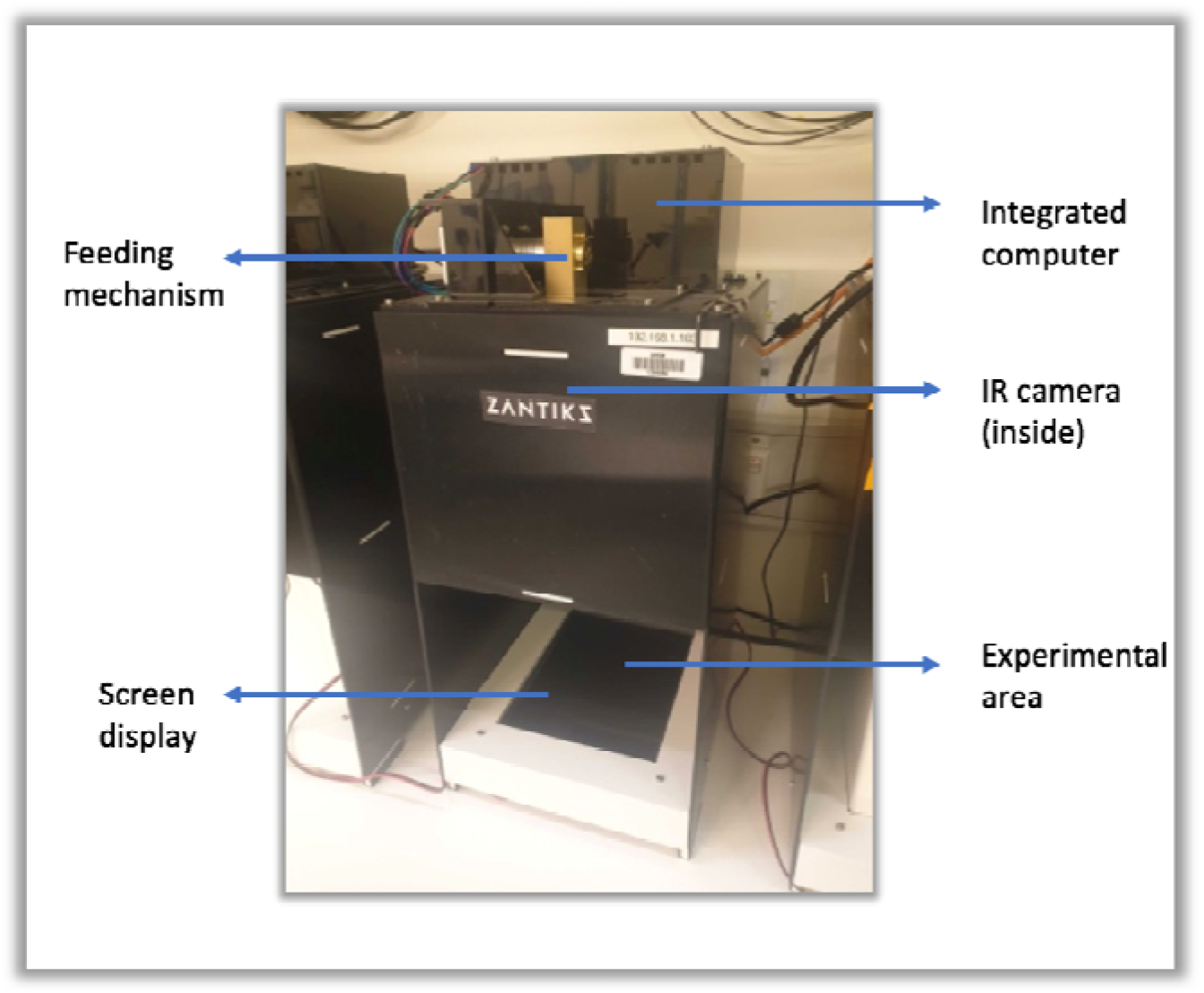
Zantiks AD unit. Fully automated experimental box with tracking (IR camera), recording (integrated computer interfaced via console, see Supplementary Figure 2E below) among other capabilities with an open compartment where the assay was conducted, with a screen that holds the tank with experimental subjects during trials.

**Supplementary Figure 2.**
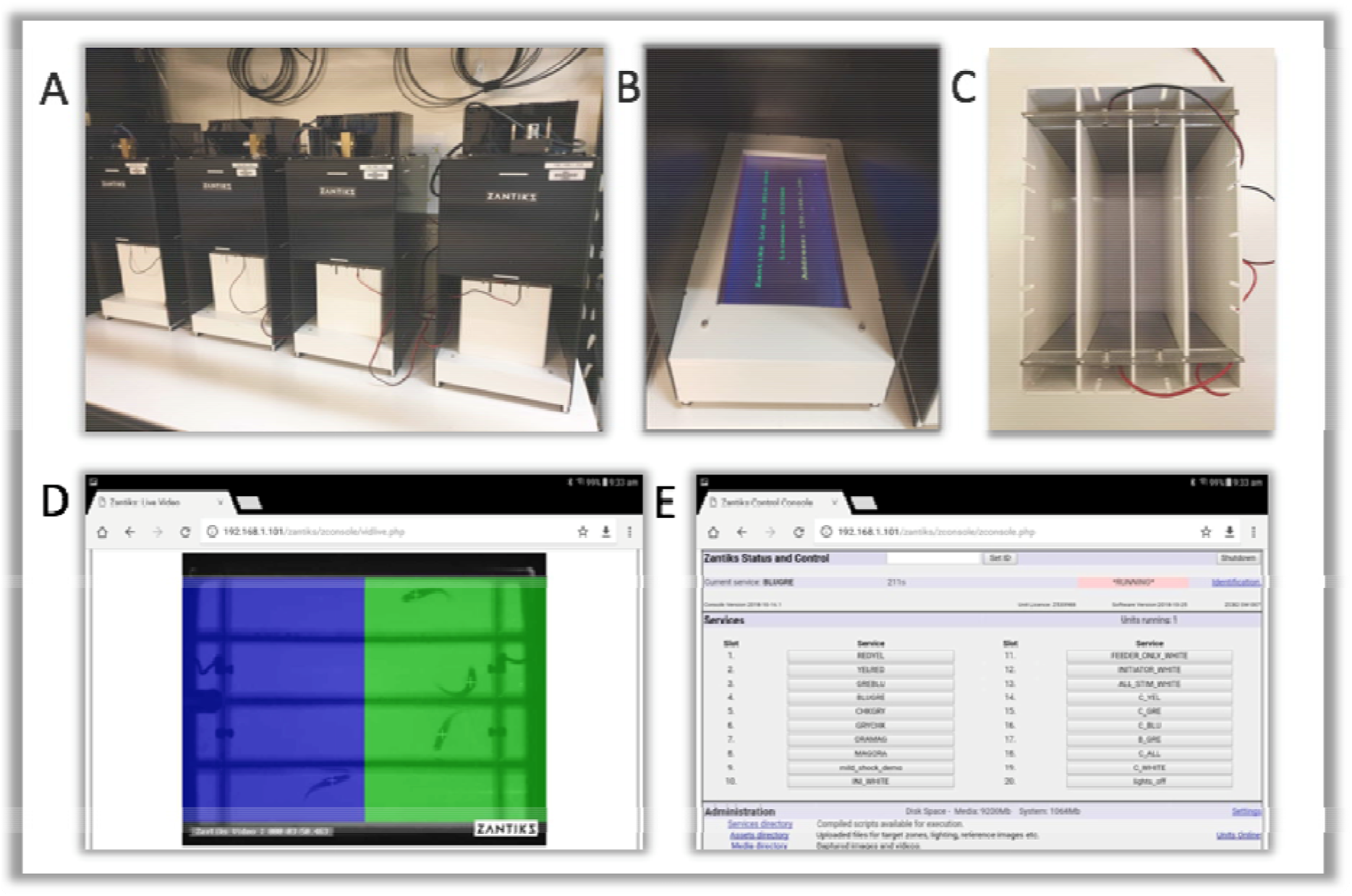
Automated conditioning setup. A: All four Zantiks AD experimental units. B: Stimulus screen programmed to present a variety of colours, patterns and images. C: Tank organised for aversive experiment with 4 lanes and 2 mild electric shock plates. D: View of example assay, depicting fish tracking and overlay of perimeters (CS zones). E: Control console for ease of interface with the AD units.

**Supplementary Figure 3.**
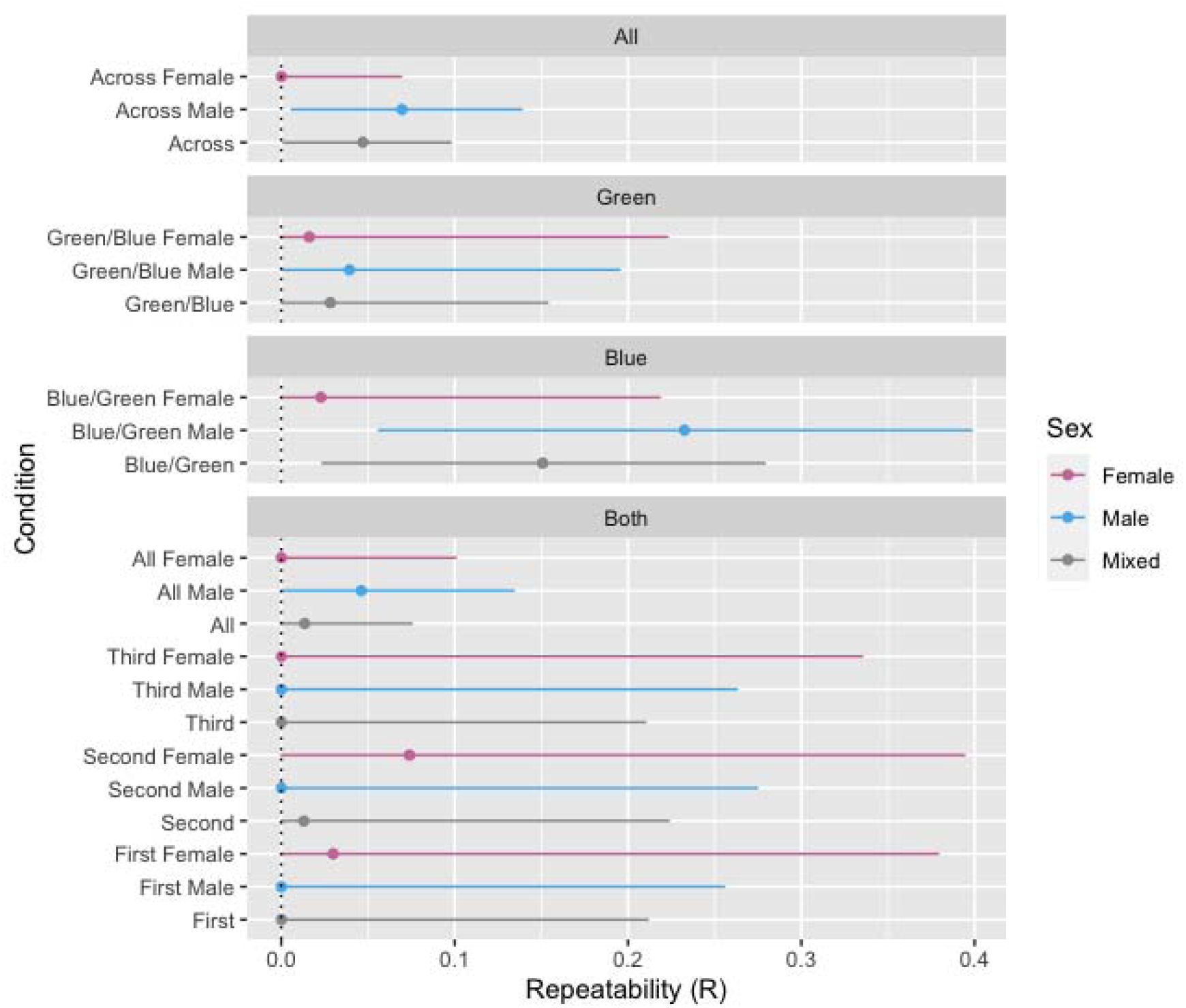
The repeatability (*R*) of aversive learning with 95% CIs in females, males and both sexes together (mixed). The segments are from top to bottom: Across all conditions, the ‘Green/Blue’ condition, the ‘Blue/Green’ condition, both ‘Green/Blue’ and ‘Blue/Green’ combined. In the bottom segments, the conditions are split into four measurement sets: ‘First’, the first measurements of both ‘Green/Blue’ and ‘Blue/Green’ (set of two measurements), ‘Second’, the second measurements of both ‘Green/Blue’ and ‘Blue/Green’, Third, the third measurements of both ‘Green/Blue’ and ‘Blue/Green’, ‘All’, all measurements sets combined (total of 6 measurements).

**Supplementary Figure 4.**
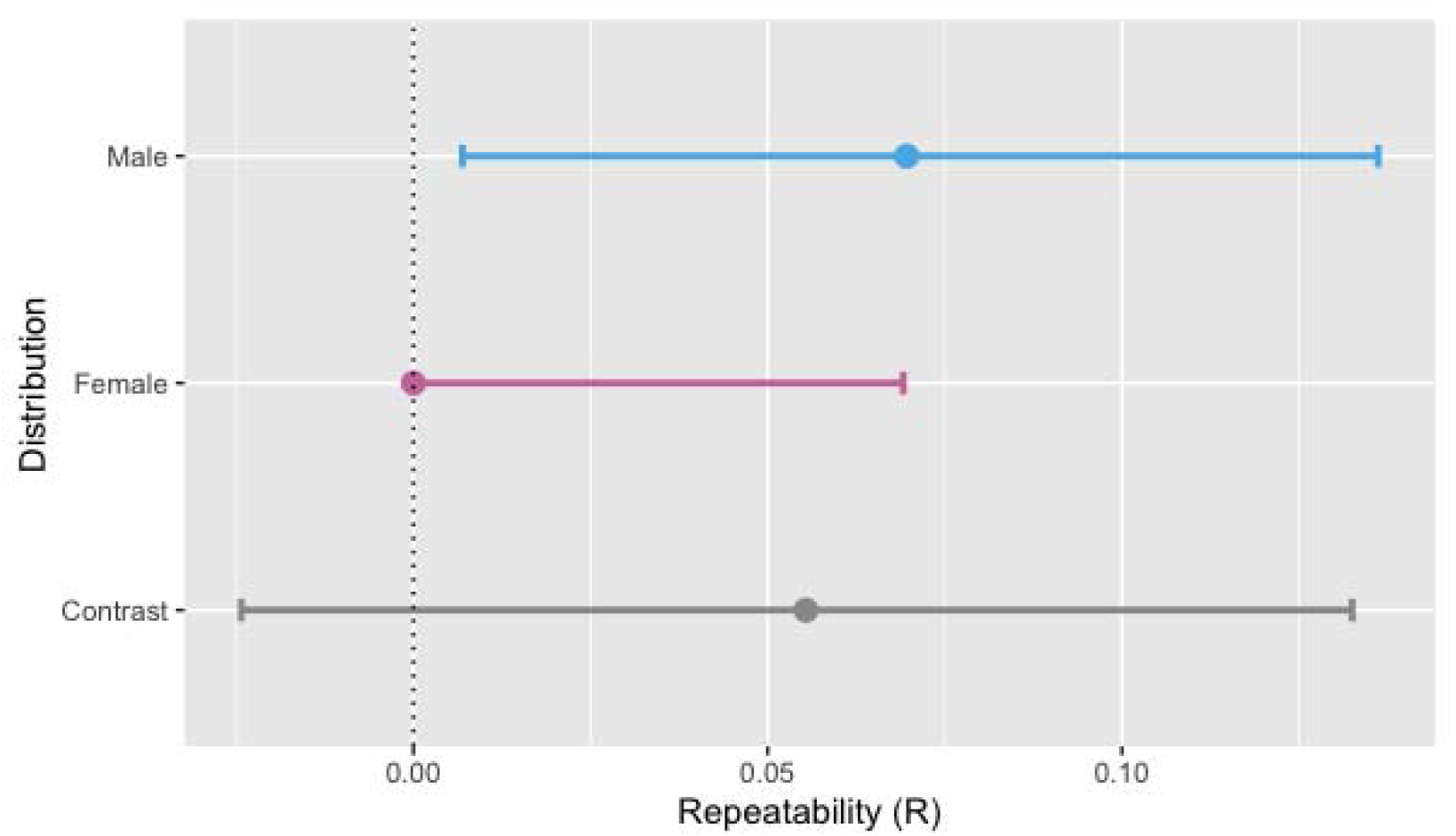
Male and female zebrafish contrast analysis of repeatability estimates in across conditions trials. Male and females differ in the repeatability bootstrap distribution, however, the contrast analysis indicates by way of the distributions overlapping zero that males and females do not significantly differ in repeatability.

**Supplementary Figure 5.**
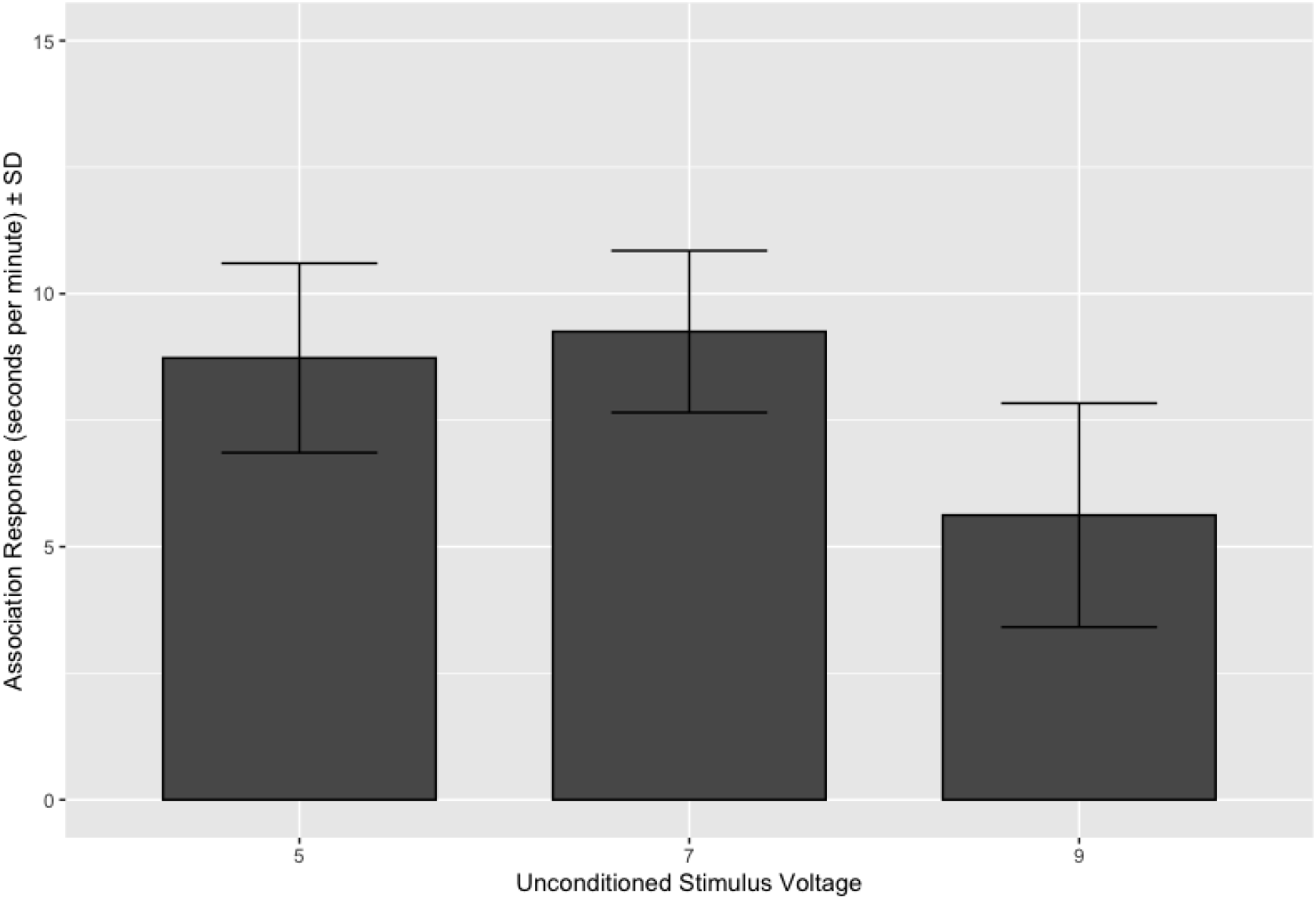
Zebrafish learning performance in three voltage settings during pilot experiment with standard deviation. Mean difference in CS+ avoidance between baseline and probe phase in seconds per minute in five, seven and nine volts.

**Supplementary Figure 6.**
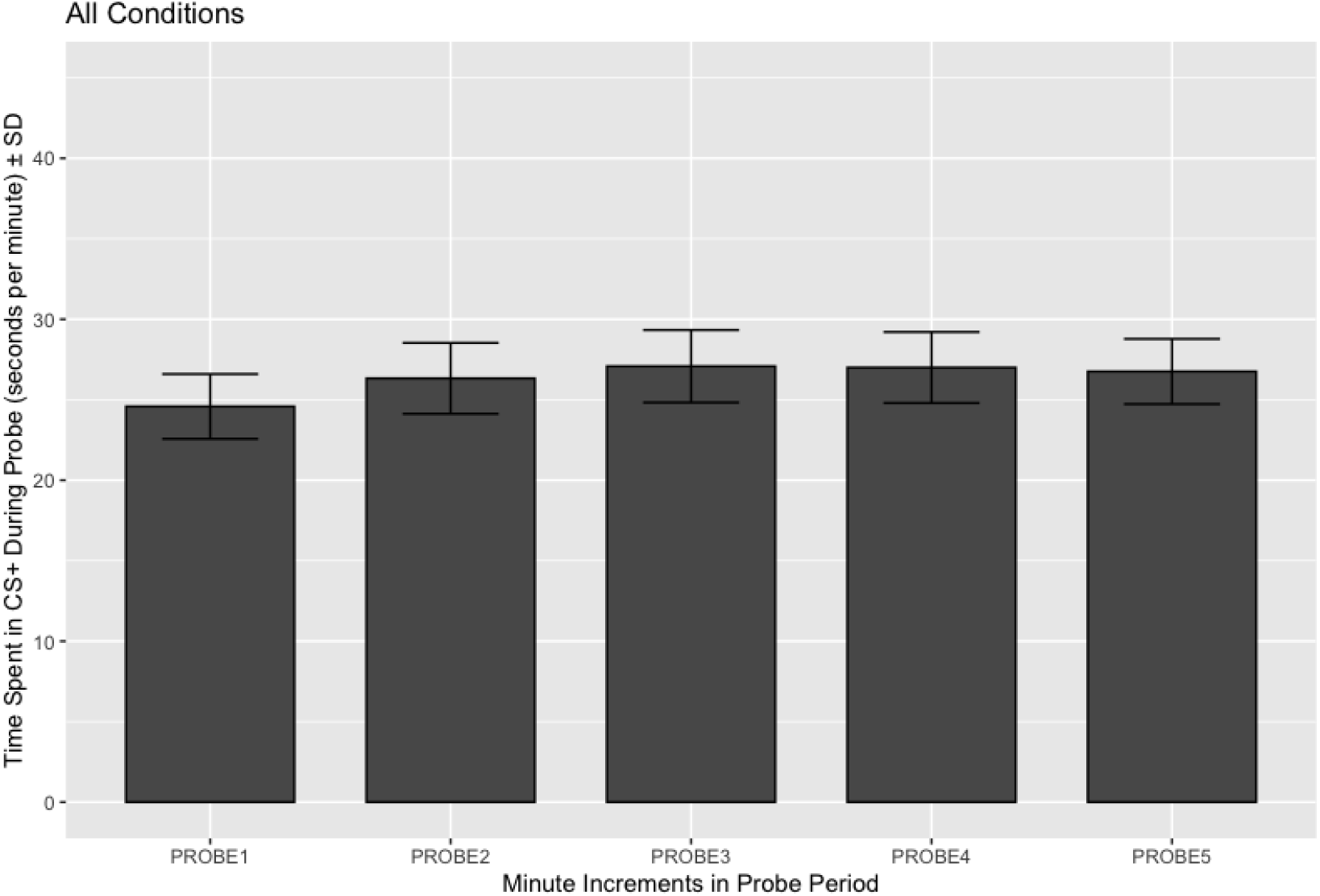
Zebrafish learning performance across all conditions in the probe period with standard deviation. Avoidance of the CS+ is shown separately for each minute of the probe.

**Supplementary Figure 7.**
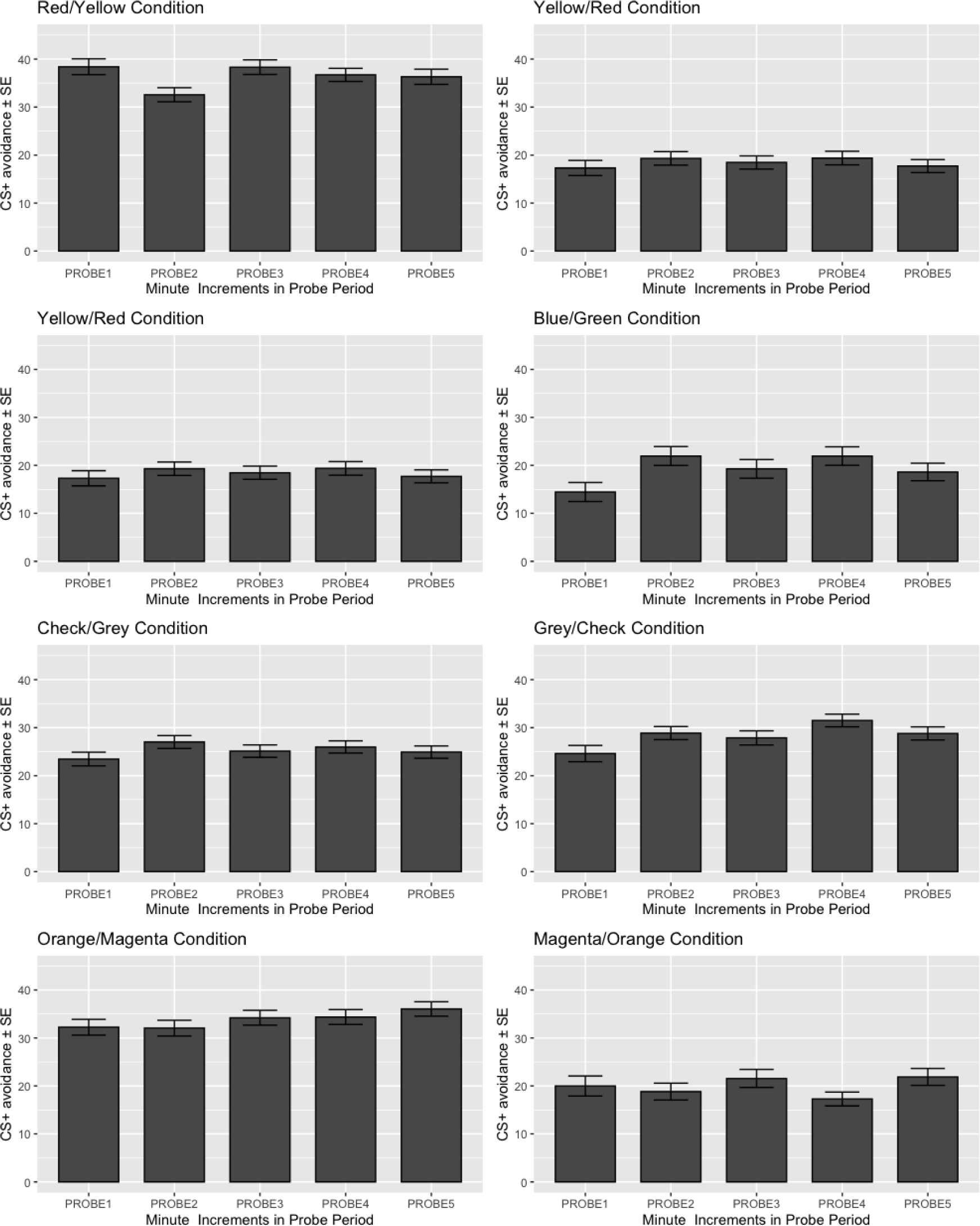
Zebrafish learning performance in each condition during the probe period with standard error. Avoidance of the CS+ is shown separately for each minute of the probe.

